# A Tensegrity-Graph Optimization Framework for Estimating Intracellular Stiffness

**DOI:** 10.1101/2021.04.04.438371

**Authors:** Raktim Bhattacharya

## Abstract

In this paper we present a formulation for estimating intracellular stiffness using tensegrity principles. We demonstrate that the new tensegrity model, based on random graphs over dense nodes, can predict well known mechanical characteristics of epithelial and mesenchymal cells.

## 1 Introduction

Mechanobiology is emerging as a field of science that is at the interface of biology and engineering, and focusses on understanding the impact of mechanical cues on cellular biology. For example, in cancer biology the emphasis is on the mechanical properties of the tumor microenvironment such as matrix stiffness, topography, stress, and cell deformation, and their influence on cellular responses, gene expression pattern, etc., which ultimately results in influencing tumor growth and proliferation [1–4].The interaction between the mechanical network and the biochemical network, also known as mechanotransduction, is a major challenge in mechanobiology.

A key step in developing mechanotransduction models is the estimation of mechanical stress in biological systems. However, the absence of tools for mapping the forces during cancer progression, has impeded our understanding of the mechanotransduction process during this phase. Although atomic force microscopy (AFM) can measure surface forces in single cells or cell doublets, and laser microsurgery provides information about the forces carried by ablated structures, comprehensive maps of driving forces have remained elusive. Fortunately, computational modeling can complement experiments and can rigorously estimate spatial and temporal maps of forces within cells.

## 2 Methods

In this work, we look at the problem of cellular stiffness estimation from a tensegrity perspective. Motivated by previous work [5], we apply tensegrity principles to model mechanical properties of cells. Tensegrity systems are essentially a network of bars (e.g. microtubules) and cables (e.g. actin filaments). For a given topology of the network, the compressive loads on the bars and tensile loads in the cables are determined from equilibrium conditions. These directly quantify the local mechanical properties (e.g. stiffness). Thus, for tensegrity systems, we can easily characterize mechanical properties, given cell-shape and biofilament topology. This leads to the possibility of estimating these properties directly from high-resolution images. Another benefit of tensegrity is that it naturally captures the auxetic behavior of cells, which requires special attention in finite-element-modeling and needs appropriate stress strain configurations. Tensegrity models of deformable structures are also possible, and is achieved by deforming the network but not the bars. Finally, simulation of tensegrity structures can be done using multi-body dynamics, which is much faster and more accurate than conventional finite element model-based simulations [6]. Thus, the tensegrity formulation scales better with respect to numerical accuracy and computational time, resulting in better models for cellular mechanics.

### 2.1 Mathematical Representation of Tensegrity Structures

Based on the formulation presented in [7], a tensegrity structure is defined by a set of nodes, a connectivity matrix defining the network of bars and cables, and the force densities (force per unit length) of the bars and cables. Let ***n**_i_* ∈ ℝ^3^ be the position of the *i*^th^ node, and ***N*** ∈ ℝ^3×*n*^ be the nodal matrix defined by

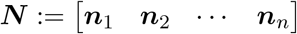

where *n* is the number of nodes in the tensegrity system. Let ***C*** ∈ ℝ^*m*×*n*^ be the connectivity matrix that defines the tensegrity system, where *m* members are defined by connecting *n* nodes. Specifically, if the *k*^th^ member is defined by connecting nodes ***n**_i_* and ***n**_j_*, then

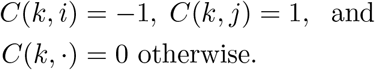

If ***σ*** ∈ ℝ^*m*^ is the vector of the force densities, the forces acting on each node due to these members are given by,

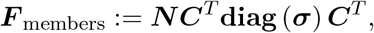

where **diag**(***σ***) represents the diagonal matrix obtained with entries of ***σ***. To impose boundary condition on the nodes, we introduce unknown reaction forces at each node ***R*** ∈ ℝ^3×*n*^. Known external forces acting at each node can also be introduced as ****F**** _external_ ∈ ℝ^3×*n*^, which can include weight of the bars and cables.

Static equilibrium is then achieved by solving for ***σ*** and ***R*** from

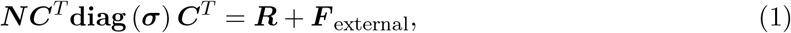

which is an underdetermined linear system of equations. Assuming existence conditions are satisfied, there are infinite solutions and an optimization problem can be formulated to arrive at a unique solution. This is discussed in the next section in the context of topology optimization.

### 2.2 Biofilament Topology from Optimization on Random Graphs

A key step in tensegrity modeling is the topology. Earlier work on tensegrity modeling of cells have resorted to polyhedrons [5, 8–10] and focuses on estimating macroscopic cellular properties. Such coarse models are unsuitable for estimating stress maps within the cell. For that we require fine grained topology that mimics the actual biofilament topology. In theory, high resolution images can be used to determine the actual biofilament topology and is an active area of research. Tools such as FiberApp [11], SOAX [12], and SIFNE [13], are being developed to determine biofilament topology from images. However, currently these tools require manual tuning and is time consuming. If the actual topology is available, it can be used to obtain the ***N*** and ***C*** matrices, and a constrained optimization can be performed to determine ***σ*** and ***R*** for a minimum stress configuration. However, if the actual topology is not available, the tensegrity topology can be over parameterized as a graph over randomly distributed points, and an optimization problem can be formulated to determine the sparsest topology that satisfies the boundary conditions and achieves minimum stress.

We next present such an optimization formulation to generate tensegrity structures. Given the cell shape and location of the nucleus (obtained from images), we distribute random nodes in the cytoplasm. The nucleus is treated as a rigid-body suspended in the cytoplasm. Although, it can also be modeled as a flexible tensegrity structure with polyhedron topology, and will be considered in our future work. Besides the nodes in the cytoplasm, we also distribute nodes on the surface of the nucleus and on the surface of the cell. Let all these nodes be represented by ****N****. The connectivity matrix ***C*** can be constructed in many ways, and can account for constraints in the biofilament topology, such as maximum length of biofilaments. An all-to-all connectivity matrix is possible, but may lead to very large number of members, and can result in computational issues. A more tractable option is to connect each node with *k* nearest neighbor [14], which is what we apply in this paper. Another option is to use techniques from random graph theory [15] to construct ***C***.

In this formulation, we account for cell mass as point masses at each node. Mass of the microtubules and actin filaments are also accounted by adding half of the mass at each of the connecting nodes. The tractile forces in the cytoskeleton are modeled using radially inward forces at each of the interior nodes. All these known forces are modeled by defining ***F*** _external_ accordingly. Finally, the cell adhesion to the ECM is modeled using unknown reaction forces ****R****. Although the variable ***R*** is defined at each node, in this formulation we only treat the components of ***R***, corresponding to boundary nodes, as unknown variables. The rest of the components of ***R*** are set to zero.

Mathematically, the optimization problem is given by

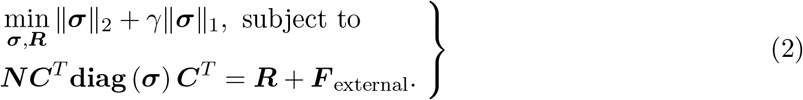

The cost function in eqn.(2) is defined by 2-norm and 1-norm of ***σ*** and represents a tradeoff between minimum stress (∥***σ***∥_2_) and minimum mass (∥***σ***∥_1_) respectively, with *γ* as a tuning parameter.

The effective topology is determined by components of ***σ***. Members with positive values of *σ_i_* are tensile and those with negative values of *σ_i_* are compressive. Therefore, the above optimization algorithmically determines the distribution of actin filaments (tensile elements) and microtubules (compressive elements), for achieving mechanical equilibrium. Furthermore, the magnitude of *σ_i_* determines the stress level of each member. Members with zero or very small ∣*σ_i_*∣ do not make significant mechanical contribution and can be removed from the topology to arrive at a lower complexity tensegrity structure.

### 2.3 Computation of Cellular Stiffness from Tensegrity Models

We next consider computation of cell stiffness from tensegrity models. Given ***N***, ***C*** and ***σ***, the stiffness matrix ***K*** can be computed as [7]

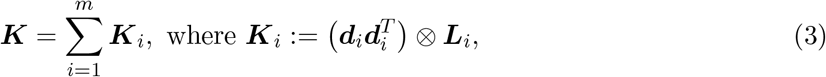

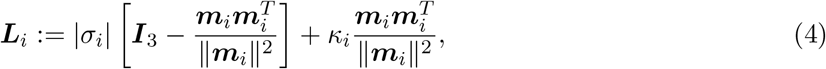

where ***d**_i_* is the *i*^th^ column of ***C**^T^*, ***m**_i_* is the *i*^th^ column of ***NC**^T^*, *κ_i_* = *E_i_A_i_/l_i_* is the material stiffness of the member, *E_i_* is the Young’s modulus, *A_i_* is the cross-section area, *l_i_* is the length of the member, *σ_i_* is the *i*^th^ of ***σ***, and ***I***_3_ is 3 × 3 identity matrix. The data for micro-tubules and actin filaments are shown in table 1.

**Table 1:** Material properties and dimension of biofilaments used in computation of cellular stiffness. [16]

|  | Microtubule | Actin Filament |
| --- | --- | --- |
| Young's Modulus | 1.2 GPa | 0.14 GPa |
| Radius | 15 nm | 3 nm |

As the cell deforms, ***N***, ***C*** and optimal ***σ*** will also vary with time, resulting in a time varying stiffness matrix. The spatio-temporal variation in the cell’s stiffness is then given by the eigen-values and the eigen-vectors of the time-varying stiffness matrix.

## 3 Results & Discussion

### 3.1 Estimation of Tensegrity Topology

We consider a circular-shaped cell with a nucleus at the center, representing epithelial cells. Fig.1a shows the over parameterized random graph with 1515 uniformly distributed nodes in a circular annulus region (obtained via acceptance-rejection sampling), 100 nodes to define the cell boundary, and 20 nodes to define the nucleus boundary. These nodes were connected using 15 nearest points. After removing members that cut across the nucleus, the topology resulted in 16033 biofilaments. To account for the cell mass, we added mass at each of the interior nodes. Mass of the nucleus was distributed over its boundary nodes. Unknown reaction forces were imposed on the nodes at the cell boundary to model the contact forces between cell and substrate, assuming a zero displacement boundary condition. Finally, we applied radially inward force towards the nucleus, at each interior node, mimicking the cell pulling on the substrate to which it is anchored. This force was arbitrarily chosen to be 15% of the nucleus weight, and distributed uniformly over the cytoplasm. A better model of the tractile force can be incorporated using available data [22]. Mechanical properties used in the optimization are given in table 2.

**Figure 1:**
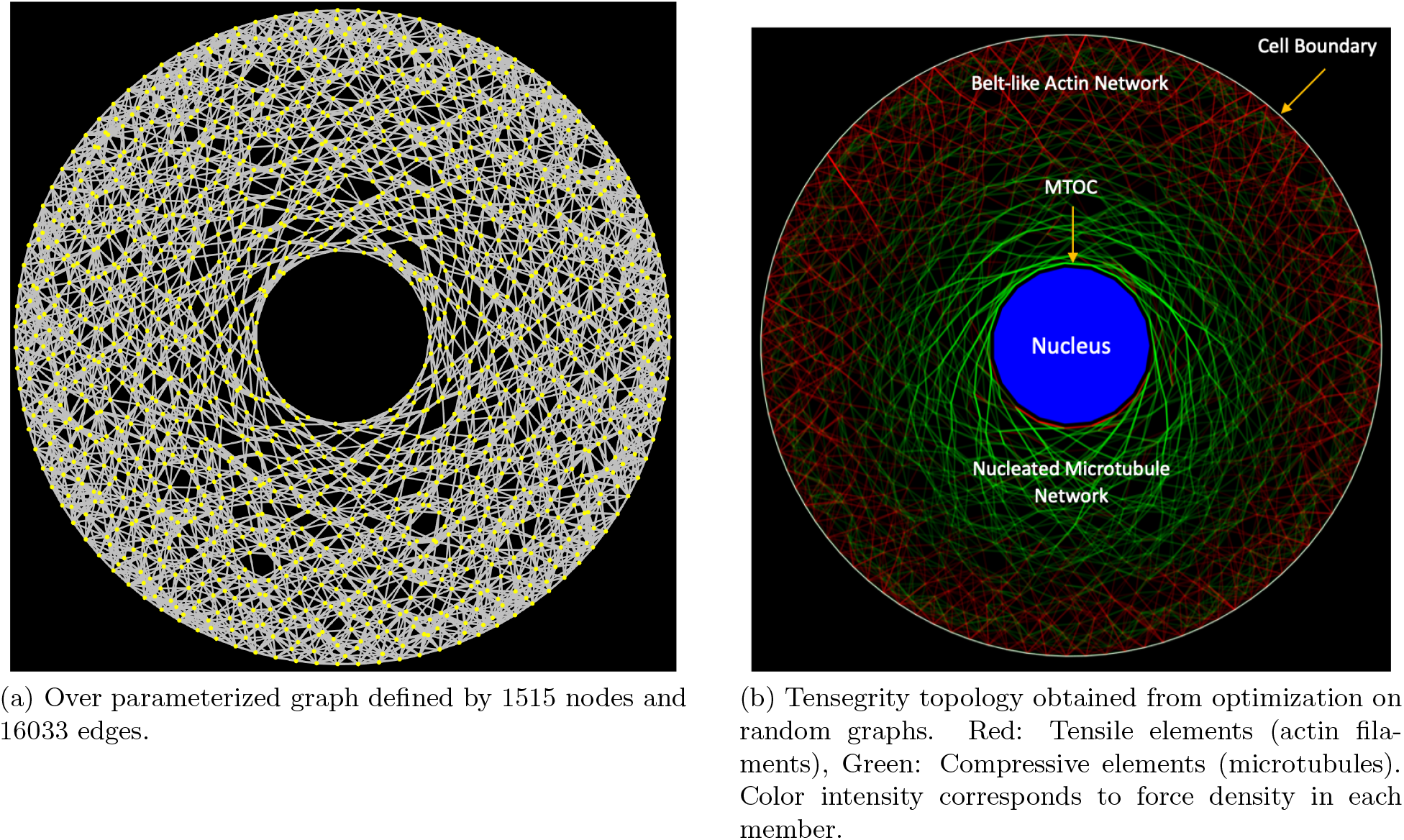
Tensegrity topology estimation from random graph optimization.

**Table 2:** Mechanical properties used in the topology estimation [16, 23, 24].

|  |  |
| --- | --- |
| Cell radius | 15.4 $\mu\text{m}$ |
| Nucleus radius | 3.5 $\mu\text{m}$ |
| Density of Cytoplasm | $1.0 \times 10^3 \text{ kg/m}^3$ |
| Density of Nucleus | $1.96 \times 10^3 \text{ kg/m}^3$ |

For this random graph, the optimization problem has 48100 variables with 18863 equality constraints. It took 43.62 seconds to solve in MATLAB, using CVX [25] and SDPT3 [26], in a MacBook Pro. The optimal solution for ***σ*** resulted in the tensegrity structure shown in fig.1b. The green lines correspond to compressive members, and the red lines correspond to tensile members. The intensity of the color indicates the magnitude of *σ_i_*, the force density of the *i*^th^ member. Thus, filaments with low force density are shown with lighter shades of red or green. From fig.1b, we make the following important observations.

The compressive elements are nucleated. The compressive elements that carry high loads originate from a common point, suggesting a microtubule-organizing center (MTOC). From a material failure perspective, these high load bearing compressive elements are expected to be thicker than tensile elements with the same load. It is also interesting to note that the MTOC is adjacent to the nucleus, just like the centrosome. We also observe the compressive elements start out from the MTOC and stretch out to the edges of the cell. All these are characteristics of the microtubule network found in cells, and our optimization-based formulation is able to recover it.

The tensile elements are shown in red and are concentrated on the outer periphery. This is similar to the actin cytoskeleton, which is commonly anchored to regions of cell-cell contact called adherens junctions. In epithelial cells, such as the model considered here, these junctions form a continuous belt-like structure (called an adhesion belt) around each cell, which is also observed here. Therefore, the network of tensile biofilaments, obtained from the optimization, closely mimic the actin networks commonly found in cells.

### 3.2 Estimation of Stress and Stiffness from Images

Using the tensegrity modeling framework presented above, we next show how we can directly quantify mechanical stress in cells by observing changes in cell shapes and their internal topology. This work is related to the visual force sensing technique common in video force microscopy (VFM) [27]. Our work is similar to VFM, as forces are determined from images using a computational model. However, VFM is based on finite-element discretization of the cells, and we pursue a tensegrity-based approach here.

Here we apply tensegrity-cell models to predict changes in the cell stiffness and forces in the cell-substrate boundary, during epithelial to mesenchymal transition (EMT). It is widely known that EMT is the key driver of cancer progression, and predicting cell stiffness and cell-ECM interaction is important for developing corresponding mechanotranduction models.

Fig.3 shows the variation in the force-density (∥***σ***\*∥_2_) in the tensegrity structure, and the reaction force (∥**R**\*∥_2_) on the ECM, as the cells undergo EMT. During EMT, the cell changes from a spherical shape (epithelial) to a spindle-like shape (mesenchymal).

From the images of the cells (as shown in fig.2a), we approximated the cell-shape with ellipses and determined the tensegrity topology using the optimization described above. Synthesis of tensegrity models, assuming static equilibrium is justified as we assume creep (quasi-static) deformation [28, 29]. As the cell deforms during EMT, the optimization is solved repeatedly to arrive at time varying ***σ***\* and ***R***\*. Repeatedly solving for ***σ***\* also allows for new topologies to be created, as it occurs in cells during EMT. Thus, this results in a spatio-temporal characterization of ∥***σ***^*^∥_2_ and ∥***R***^*^∥_2_. Higher resolution is achieved by increasing the number of bars and cables in the network, at increased computational cost.

Fig.2b plots ∥***σ***^*^∥_2_ and ∥***R***^*^∥_2_ as the cell undergoes EMT. Since, cell stiffness is proportional to ****σ****, and the stress on the ECM is proportional to the reaction forces ***R*** at the cell boundary, trends in ∥***σ***^*^∥_2_ and ∥***R***^*^∥_2_ qualitatively present the trend in cell stiffness and the stress on the ECM. We observe that our tensegrity-cell model is able to correctly predict macroscopic-mechanical properties of cells during EMT. Specifically, we make the following key observations.

It is well known that epithelial cells undergo cell softening as they acquire malignant features. This is exactly what we observe from the model. From fig.2b (top panel), we see that the model predicts a loss in cell-stiffness, as the cell transforms from circular shape (epithelial) to a spindle-like shape (mesenchymal).

**Figure 2:**
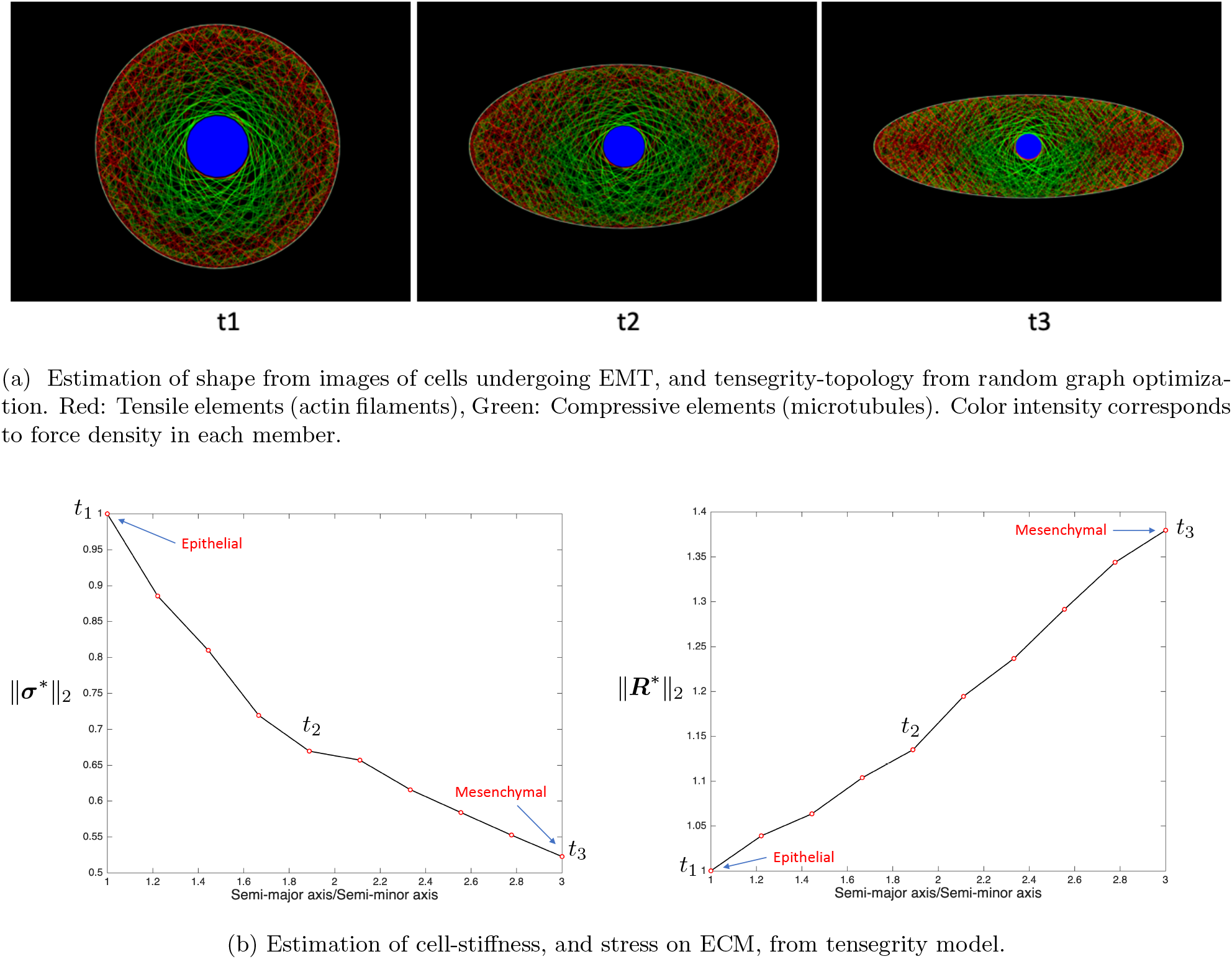
Estimation of cellular stress from images.

It is also known that during EMT transitions the ECM stiffens, which results in higher-stress at the cell-ECM junction. This is also predicted by our model. From fig.2b (bottom panel), we see that the total stress at the boundary increases as the cell transforms from circular to spindle shape.

From the tensegrity topology in fig.2a, for mesenchymal cells (spindle-shaped), we see clusters of tensile members (or actin filament network) at the extreme ends of the semi-major axis. We also observe in the tensegrity model that the actin density decreases with the distance from the leading edge, and thus it captures the well known depolymerization effects during EMT. One of the extreme ends become the leading edge of motile cells. This is achieved by forming lamellipodia, which promotes motility.

The plots for ∥***σ***^*^∥_2_ and ∥***R***^*^∥_2_ in fig.3 are normalized with respect to the values from the epithelial cell at *t*_1_, which are 9.6825 × 10^−12^ N/m and 3.1197 × 10^−13^ N respectively.

**Figure 3:**
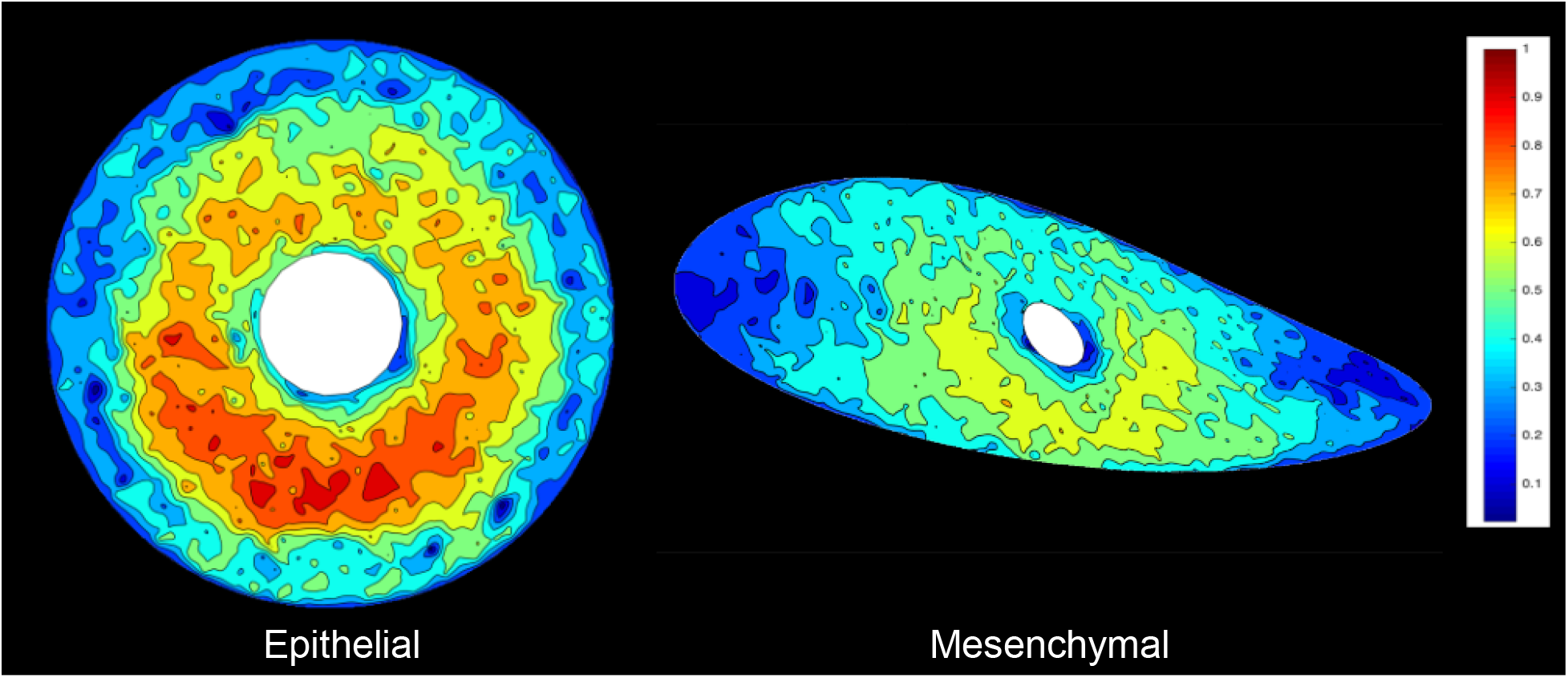
Tensegrity model-based estimation of cell stiffness in epithelial and mesenchymal cells. The color map indicates the stiffness values, normalized by 661.09 N/m, which is the maximum stiffness in the epithelial cell.

### 3.3 Cellular Stiffness Distribution from Tensegrity Models

Using eqn.(4), we compute the stiffness matrix ***K*** associated with the tensegrity cell model. Fig.3 is obtained via weighted summation of the eigen-vectors of ***K***, with the eigen-values of ***K*** as the weights. This results in the total stiffness at each point. While fig.2b (top) shows the total cellular stiffness, fig.3 shows the distribution of the stiffness for epithelial and mesenchymal cells. The color axis has the same range for both. In the plot, the stiffness values have been normalized by the maximum stiffness in the epithelial cell, which is 661.09 N/m. As expected [30], the circular or the epithelial cell, has higher stiffness in comparison with the spindle-shaped or mesenchymal cell. We make the following key observations with respect to the predictions from the tensegrity model.

Fig.3 shows the model predicted stiffness distribution in epithelial and mesenchymal cells. We observe that the stiffness of epithelial cells is higher in the central areas over cell nuclei than in peripheral areas, which matches experimental observations reported in the literature [31]. Fig.3 shows the model predicted stiffness distribution in mesenchymal cells. We observe that the extreme ends of the semi-major axis have reduced rigidity, and are thus more flexible than the epithelial cells. This matches what is observed in mesenchymal cells where the rigidity of the lamellipodia decreases, which facilitates contraction of the actin/myosin II network at the lamellipodium/cell body boundary [32, 33]. In addition, the mesenchymal cells are generally softer and display lower intrinsic variability in cell stiffness than non-malignant epithelial cells [34]. We also observe in mesenchymal cells that there is a reduction in stiffness around the nucleus. This supports the fact that mesenchymal cells have softer nucleus to increase cell motility and migration [35, 36]. Thus, the cellular tensegrity model is able to capture some key mechanobiological aspects of EMT, which is extremely encouraging.

